# An interaction network of the human SEPT9 established by quantitative mass spectrometry reveals an interplay with myosin motors

**DOI:** 10.1101/395996

**Authors:** Matthias Hecht, Reinhild Rösler, Sebastian Wiese, Nils Johnsson, Thomas Gronemeyer

**Author notes:** corresponding author: PD Dr. Thomas Gronemeyer, Ulm University, Institute of Molecular Genetics and Cell Biology, James Franck Ring N27, 89081 Ulm, Germany.

## Abstract

**Summary:** Phenotypic variations of an organism or a single cell often arise from alterations of protein interaction networks. We provide here an interaction network of the human SEPT9, an important component of the human septin cytoskeleton.

**Abstract:** Experimentally deciphering and understanding the interaction network of a particular protein provides often evidence for so far unknown functions. For the septins, a class of cytoskeletal proteins, targeted high-throughput approaches that aim at systematically deciphering interaction partners have not yet been performed. Septins regulate the organization of the acin cytoskeleton, vesicle transport and fusion, chromosome alignment-and segregation, and cytokinesis. SEPT9 is part of the core septin hetero-octamer in human cells which is composed of SEPT2, SEPT6, SEPT7, and SEPT9. SEPT9 has been linked to a variety of intracellular functions as well as to diseases and diverse types of cancer. We applied a quantitative proteomics approach to establish an interactome of SEPT9 in human fibroblast cells. We identified among others so far unknown interaction partners from the myosin family and could provide evidence that SEPT9 participates in vesicle transport from and to the plasma membrane as well as in the attachment of actin stress fibers to cellular adhesions.

## Introduction

The septins are often named the fourth component of the cytoskeleton and gained a growing attention over the past few years [1]. However, molecular mechanisms on septin assembly are much less understood than those of microtubules, actin filaments and intermediate filaments [2].

The mammalian genome encodes thirteen different septins (SEPT1-SEPT12, SEPT14) [3], of which SEPT2, SEPT7 and SEPT9 are nearly ubiquitously expressed, while SEPT1, SEPT3, SEPT12, and SEPT14 are tissue-specific. The remaining septins are expressed widely, but are not present in all tissues [4]. All septin subunits can be sorted into four subgroups based on sequence homology, namely the SEPT2, SEPT3, SEPT6 and SEPT7 subgroup [5].

In mammalian cells, septins regulate the organization of the cytoskeleton, vesicle transport and fusion, chromosome alignment, segregation, and cytokinesis [6–10]. Septins cross-link and bend actin filaments into functional structures such as contractile rings in cytokinesis or stress fibers in filopodia and lamellipodia during cell migration [11]. Mammalian SEPT2/6/7/9 complexes do also interact directly with microtubules [12].

The basic septin oligomer in mammalian cells is a hetero-octamer composed of the SEPT2, SEPT6, SEPT7, and SEPT9 at a stoichiometry of 2:2:2:2 (9-7-6-2-2-6-7-9). Septin oligomers polymerize into higher ordered structures such as rings, filaments, and gauzes by end-to-end and lateral joining. The assembly of septins does not only depend on their biochemical properties and posttranslational modifications, but also on intracellular structures and interaction partners.

Given the range of cellular functions associated with septins, it is not surprising that a variety of diseases are linked to loss or gain of septin functions. Abnormalities in septin expression are linked to male infertility, hereditary neuralgic amyotrophy, neurodevelopmental disorders, Alzheimer, and Parkinson [4,13–15].

Especially SEPT9 has been linked to a variety of diseases and to diverse types of cancer including prostate, breast and colon cancer [16–19]. This member of the septin family is characterized by a number of splice variations. The SEPT9 locus contains 13 exons and encodes at least nine isoforms (according to NCBI Reference Sequences, Gene ID: 10801). The SEPT9 gene has unique characteristics as it as been shown to act as an oncogene, although tumor suppressive properties have been reported as well [20].

SEPT9 is linked to several intracellular processes. SEPT9 binds and bundles microtubules [12] and plays an important role in cytokinesis by mediating the localization of the vesicle-tethering exocyst complex to the midbody [7]. It interacts furthermore with F-actin and functions as a stress fiber cross-linking protein [21]. However, for the septins in general and SEPT9 in particular, no targeted high throughput aproaches aiming at systematically deciphering interaction partners have been conducted that could nourish the beforehand described systematic studies. Nowadays it is established that proteins act in networks and phenotypic variations of an organism or a single cell often arise from alterations of these networks [22]. Thus, experimentally deciphering and understanding the network environment of a protein often complements classic, systematic approaches [23].

We addressed this issue by applying a quantitative proteomics approach to establish an interactome of SEPT9 in human fibroblast cells. We could identify new interaction partners from the myosin familiy and could provide evidence that SEPT9 participates in vesicle transport from and to the plasma membrane as well as in the attachment of actin stress fibers to cellular adhesions.

## Results and Discussion

We employed a SILAC based workflow coupled to affinity purification (AP-MS) [24] for the investigation of the interactome of SEPT9_i1 in human 1306 fibroblast cells. First, we constructed a TAP-SEPT9 expressing cell line and a GFP-SET9 expressing cell line that served as the control cell line. Both fusion proteins were expressed under the control of a CMV promoter. However, when using tagged proteins in an overexpression system, the correct localization of the tagged protein has to be ensured. We therefore investigated first if consistent expression of tagged SEPT9 interferes with the endogenous septin cytoskeleton by immunostaining the ubiquitously expresssed SEPT7. Both GFP-SEPT9 and TAP-SEPT9 colocalized in filamentous structures with the endogenous septin cytoskeleton (Fig. 1A). A portion of the tagged SEPT9 was localized to the nucleus. Whether this represents an artifact could not be verified but as we used exclusively the cytoplasmic fraction for subsequent MS analysis this issue was not investigated further. It was already reported that overexpression of GFP-SEPT9_i1 led to pronounced localization to the nucleus in breast cancer cells [16].

**Figure 1.**
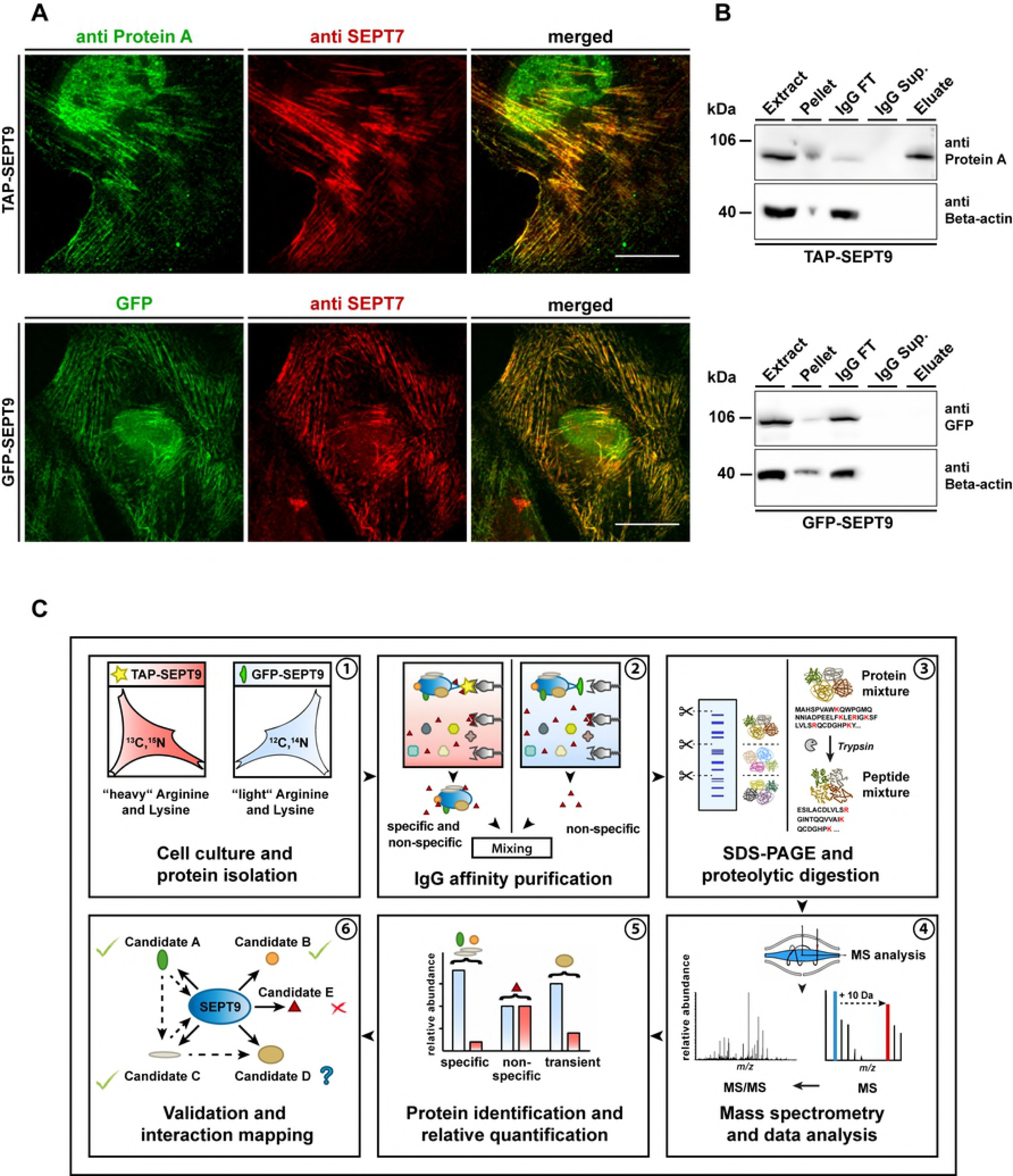
Cell lines and workflow for AP-MS of SEPT9 complexes. **A)** Characterization of the used cell lines. The colocalization of GFP-SEPT9 (upper panel) and TAP-SEPT9 (lower panel) with the endogenous septin cytoskeleton was probed by immunostaining of SEPT7. TAP-SEPT9 was visualized via an anti-ProteinA antibody. The scale bar represents 20 µM. **B)** Western blot analysis of the consecutive steps of the AP. Samples of the cell extract, pellet, supernatand of the beads after coupling (IgG sup.), washing step (IgG wash) and the eluate were separated by SDS-PAGE and Western blot was performed using an anti-ProteinA or anti-GFP antibody, repectively. **C)** Workflow of the performed “mixing after purification” AP approach with subsequent MS analysis.

For the identification of septin-interacting proteins, SEPT9 complexes were subsequently affinity-purified from the cytoplasmic fraction in a mixing after purification (MAP) approach that allows also the identification of transient interactors [25]. The TAP-SEPT9 expressing cell line was previously labeled with “heavy” SILAC amino acids. Equal volumes of eluates were combined prior to SDS-PAGE and subsequent LC/MS analysis. Affinity purifications were monitored by Western blot analysis of representative samples taken during the purification process (Fig. 1B). As expected, TAP-SEPT9 could be detected in the elution from the affinity matrix whereas GSP-SEPT9 could not. The whole AP-MS workflow is summarized in figure 1C.

We performed three independent AP-MS purifications. For each identified protein we calculated a “heavy over light” (H/L) ratio and significance B values (SigB) [26]. The LOG_2_(H/L) reflects the enrichment of a protein in the TAP-SEPT9 purification (H) in comparison to the GFP-SEPT9 purification (L). SigB represents an outlier significance score for log protein ratios which takes the peptide intensities into account.

We considered a protein in a replicate as significantly enriched at a LOG_2_(H/L)>2. The threshold value for significance according to SigB was 0.05.

Two categories of SEPT9 interacting proteins were initially defined: Class I interactors had a LOG_2_(H/L)>2, were significant according to SigB and were identified in at least two of the three replicates. Only 8 proteins could be assigned to this category. Many *bona fide* interactors with a high LOG_2_(H/L) ratio were only identified in one replicate. However, a replicate represents a frozen snapshot of both stable and transient interactions with the bait protein in a heterogenous cell population. We consequently definied a second category of interactors with loosened criteria compared to Class I. Class II interactors had a LOG_2_(H/L)>2, were significant according to SigB or identified in at least two replicates. In this category 28 candidates are listed and it accomodates also *bona fide* transient interactors. All proteins identified as Class I and Class II interactors including their LOG_2_(H/L)>2 ratios are summarized in Table 1.

**Table 1.** List of all indentified specific SEPT9 interactors (Class I and Class II) and manually curated Class III candidates.

|  | UniProt ID | Protein name | Gene name | AP1<br>LOG<sub>2</sub>(H/L) | AP2<br>LOG<sub>2</sub>(H/L) | AP3<br>LOG<sub>2</sub>(H/L) |
| --- | --- | --- | --- | --- | --- | --- |
| Class I | P08670 | Vimentin | *VIM* | 5.061 | 2.741 | 0.107 |
|  | O15231 | Zinc finger protein 185 | *ZNF185* | 5.600 | 2.710 | 1.282 |
|  | A6NMH6 | Septin-8 | *SEPT8* | 7.346 | - | 3.568 |
|  | Q15019 | Septin-2 | *SEPT2* | 2.167 | 2.595 | 1.856 |
|  | E7ES33 | Septin-7 | *SEPT7* | 2.639 | 4.887 | 1.778 |
|  | D6RGI3 | Septin-11 | *SEPT11* | 2.901 | 3.891 | 2.060 |
|  | D6RGI3 | Aminopeptidase N | *ANPEP* | 3.029 | 2.185 | - |
|  | Q8WWI1 | LIM domain only protein 7 | *LMO7* | - | 2.225 | 3.794 |
| Class II | E9PDF6 | Unconventional myosin-Ib | *MYO1B* | 4.846 | 2.220 | 0.623 |
|  | E7EPZ9 | Tenascin-X | *TNXB* | 7.652 | - | - |
|  | P17252 | Protein kinase C alpha type | *PRKCA* | 3.279 | - | - |
|  | Q9UBI6 | Guanine nucleotide-binding protein G(I)/G(S)/G(O) subunit gamma-12 | *GNG12* | 3.828 | - | - |
|  | P10301 | Ras-related protein R-Ras | *RRAS* | 6.174 | - | - |
|  | Q96FN4 | Copine-2 | *CPNE2* | 3.924 | - | - |
|  | E9PNW4 | CD59 glycoprotein | *CD59* | 5.288 | - | 0.635 |
|  | Q6YHK3 | CD109 antigen | *CD109* | 6.287 | - | - |
|  | Q6XZB0 | Lipase member I | *LIPI* | 3.524 | - | - |
|  | P05362 | Intercellular adhesion molecule 1 | *ICAM1* | 4.501 | - | - |
|  | H3BNE1 | Synaptosomal-associated protein 23 | *SNAP23* | 4.958 | - | - |
|  | Q86WV6 | Stimulator of interferon genes protein | *TMEM173* | 4.491 | - | - |
|  | O43707 | Alpha-actinin-4 | *ACTN4* | 2.456 | 2.004 | 0.377 |
|  | H0Y7A7 | Calmodulin | *CALM2* | 2.538 | 2.054 | 1.017 |
|  | P08754 | Guanine nucleotide-binding protein G(k) subunit alpha | *GNAI3* | 3.005 | - | - |
|  | F8W7R3 | Fanconi anemia group I protein | *FANCI* | 3.075 | - | - |
|  | Q15283 | Ras GTPase-activating protein 2 | *RASA2* | 3.116 | - | - |
|  | Q9Y2F5 | Little elongation complex subunit 1 | *ICE1* | - | - | 6.978 |
|  | Q15746 | Myosin light chain kinase | *MYLK* | - | 4.389 | 1.663 |
|  | Q9P2N5 | RNA-binding protein 27 | *RBM27* | - | 6.739 | 0.250 |
|  | Q9BX40 | Protein LSM14 homolog B | *LSM14B* | - | 3.080 | 1.798 |
|  | Q9BX40 | Protein LSM14 homolog A | *LSM14A* | - | 3.078 | 1.797 |
|  | Q03135 | Caveolin-1;Caveolin | *CAV1* | 1.941 | 3.644 | - |
|  | E9PE29 | Zinc finger protein 106 | *ZNF106* | - | 3.688 | - |
|  | Q6NYC8 | Phostensin | *PPP1R18* | - | 6.145 | - |
|  | Q8IYI6 | Exocyst complex component 8 | *EXOC8* | - | - | 6.289 |
|  | Q86UE4 | Protein LYRIC | *MTDH* | - | 3.969 | - |
|  | P00505 | Aspartate aminotransferase, mitochondrial | *GOT2* | - | - | 3.541 |
| Class III | F5H6E2 | Unconventional myosin-Ic | *MYO1C* | 4.519 | 1.450 | 0.357 |
|  | F8W6L6 | Myosin-10 | *MYH10* | 2.339 | 0.585 | 0.376 |
|  | Q9UHB6 | LIM domain and actin-binding protein 1 | *LIMA1* | 2.690 | 1.281 | 1.174 |
|  | O14974 | Protein phosphatase 1 regulatory subunit 12A | *PPP1R12A (MYPT1)* | 2.987 | 0.130 | 0.486 |
|  | Q6WCQ1 | Myosin phosphatase Rho-interacting protein | *MPRIP* | 3.061 | 0.159 | 0.560 |
|  | E7EW20 | Unconventional myosin-VI | *MYO6* | 3.767 | 1.913 | - |

All interactors were sorted according to their biological function into seven categories, namely (i) cytoskeletal and cytoskeleton associated proteins, (ii) proteins involved in proliferation, differentiation and apoptosis, (iii) cell surface proteins, (iv) proteins involved in metabolism, (v) signal transduction proteins, (vi) nucleic acid associated proteins and (vii) proteins associated with the plasma membrane. Based on these categories we draw a map of SEPT9 interactors (Fig. 2).

**Figure 2.**
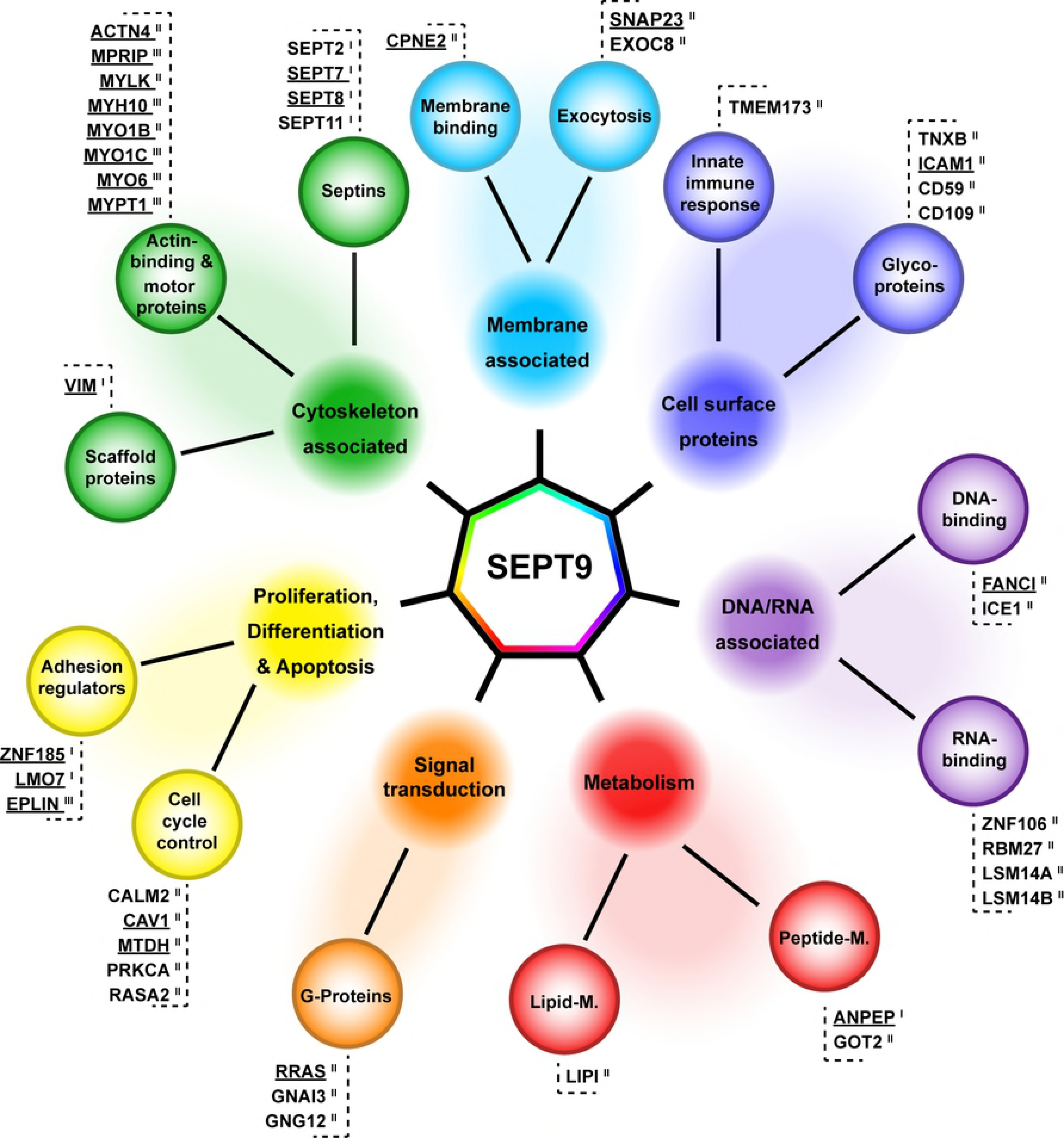
Interaction network of SEPT9 in 1306 fibroblast cells. Class I, II and III SEPT9 interactors identified by SILAC based AP-MS were categorized into 7 functional groups, (i) cytoskeleton associated proteins, (ii) proteins involved in proliferation, differentiation and apoptosis, (iii) cell surface proteins, (iv) metabolic proteins, (v) signal transduction proteins, (vi) nucleic acid associated proteins and (vii) proteins associated with the plasma membrane. Candidates validated by IF are underlined.

We subsequently defined a list of candiates for further investigation by complementary experiments:

Other septins were obvious and expected interaction candidates for SEPT9. SEPT2, SEPT7, SEPT8 and SEPT11 were found as Class I interaction partners. The interaction of SEPT9 with SEPT7 and SEPT2 was previously shown by yeast two hybrid and affinity capture experiments, respectively [27,28]. We did not detect SEPT6 as specific interactor, one of the core subunits of the canonical SEPT2-SEPT6-SEPT7 hexamer. However, the individual members within one septin subgroup can substitute for one of the others within the complex [27,28]. As SEPT6 and SEPT8 belong to the same subgroup, we conclude that in our cellular system SEPT8 is replacing SEPT6.

The intermediate filament Vimentin was identified as Class I interactor. This protein is highly expressed in fibroblasts and interacts with many proteins inside the cell [29]. Aminopeptidase N (ANPEP, CD13) is a integral membrane protein with functions in cell adhesion and processing of bioactive peptides [30]. It has a large extracellular domain and a short intracelllar C-terminal tail. We suspected that these two interaction hits were false positives and we could indeed show by IF staining of septins and Vimentin that both intracellular structures do not colocalize (Supplementary Figure 1). The antibody against ANPEP was not specific (data not shown) and we did not follow up this interaction.

Two LIM domain containing proteins, LMO7 and ZNF185 were detected as Class I SEPT9 interactors. LIM domains contain tandem zinc-finger structures and function as a modular protein-binding interfaces (reviewed in [31]). Both proteins were shown to associate with actin structures and play a role in cancer progression [32,33]. One more LIM domain containing protein, LIMA1 (herein from now on referred to as EPLIN, the product of the LIMA1 gene), was identified with a LOG_2_(H/L)>2 in one of the replicates, however below the SigB threshold. We decided to add this protein to the candidate list for further validation.

We found the presence of myosins and myosin associated proteins among the Class II candidates particularly interesting as SEPT9 competes with myosin for binding to the same sites of F-actin [21]. However, a direct interaction of SEPT9 with myosin motors has not been shown yet. Another septin, SEPT2, has been previously reported to interact directly with non-muscle myosin II [34] and thereby enables myosin II activity in interphase and dividing cells.

We screened our data set for further myosin or myosin associated proteins. We identified five proteins, MYO1C, MPRIP, MYH10 and PPP1R12A (MYPT1), with a LOG_2_(H/L)>2. However, these hits were identified each in only one replicate as SEPT9 interactor and they were not siginificant according to SigB. Several more myosin variants with a LOG_2_(H/L)<2 were identified. Among these, we picked one more candidate, MYO6, for further validation as MYO6 is the only reverse direction motor that moves towards the minus end of actin filaments [35] and as MYO6 is involved in targeted membrane transport during cytokinesis [36].

These myosin and myosin associated were grouped together with EPLIN in another category, manually curated Class III interactors (Table 1).

Besides the above mentioned Class I and the manually curated Class III interactors, we picked further candidates from each functional category of the Class II interactors (except the cell surface proteins) for validation by immunflorescence (IF) (underlined proteins in Fig. 2). Altogether, we selected SNAP23, EPLIN, ZNF185, LMO7, MYPT1, MPRIP, MYO1B, MYO1C, MYO6, MYH10, MYLK, ACTN4, CAV1, FANCI and RRAS for validation experiments.

We perfomed IF as well in the stably GFP-SEPT9 expressing cell line (Fig. 3) as in wildtype fibroblasts using an anti-SEPT7 antibody (Fig. 4). The results were consistent in both cell lines. For some tested candidates (LMO7, MYLK, ACTN4, CAV1, MYO1B), the respective antibody was unspecific and thus a defined localization was not detectable (data not shwon). FANCI and RRAS did not colocalize with the septin cytoskeleton (Supplementary Fig. 1). In the following, the IF positive candidates are discussed in more detail.

**Figure 3.**
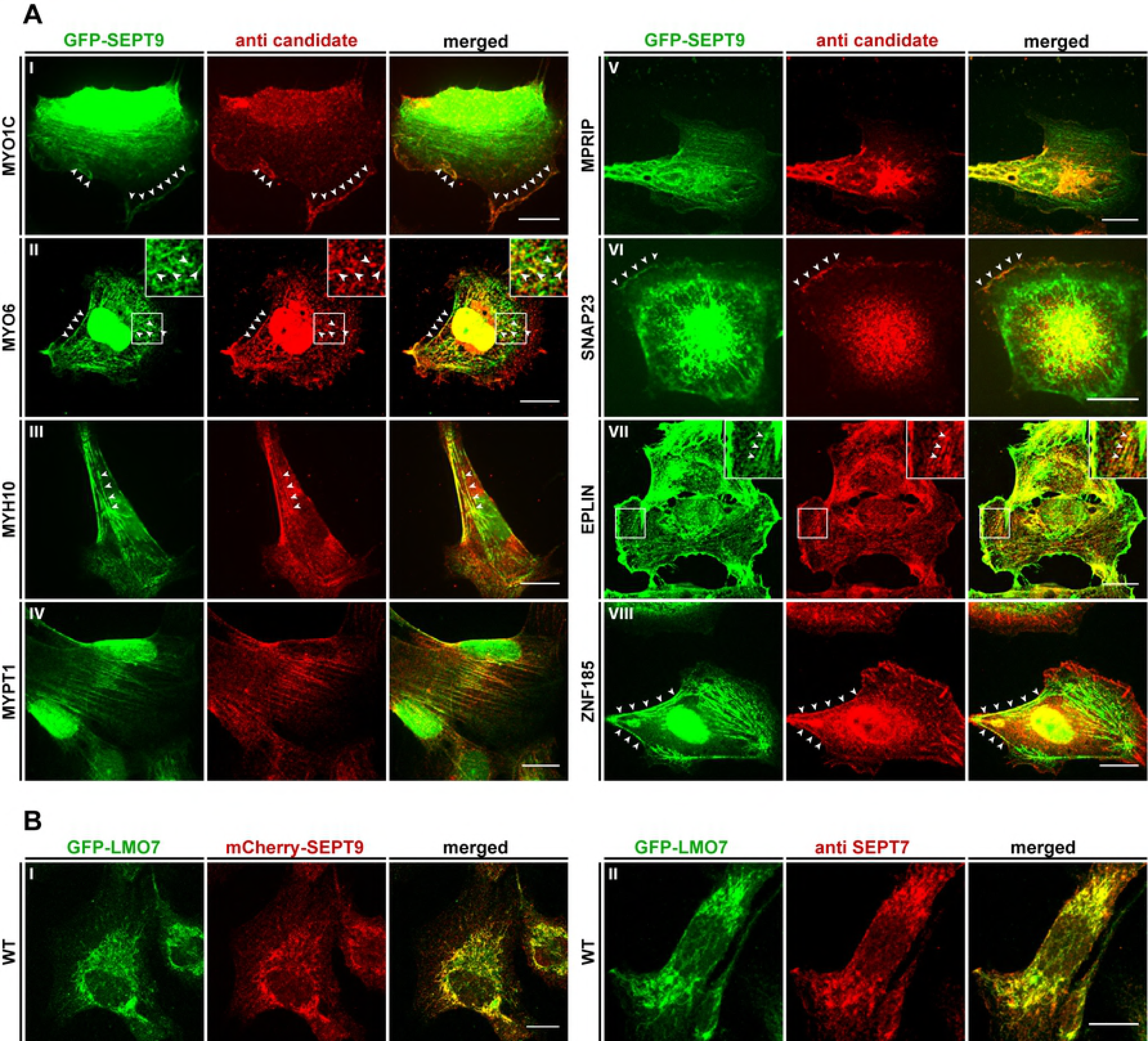
Colocalization of interaction partners with SEPT9 in GFP-SEPT9 expressing cells. **A)** The indicated candidate proteins were immunostained with a suitable primary antibody, followed by an Alexa555 coupled secondary antibody. GFP-SEPT9 was observed directly via its GFP. White arrowheads mark colocalizing structures. **B)** Colocalization of GFP-LMO7 with mCherry-SEPT9 in transiently transfected 1306 cells (left panel) and colocalization with the endogenous septin cytoskeleton by IF via an anti-SEPT7 antibody (right panel). Images were assembled from z-projections. The scale bars represent 20 µM.

**Figure 4.**
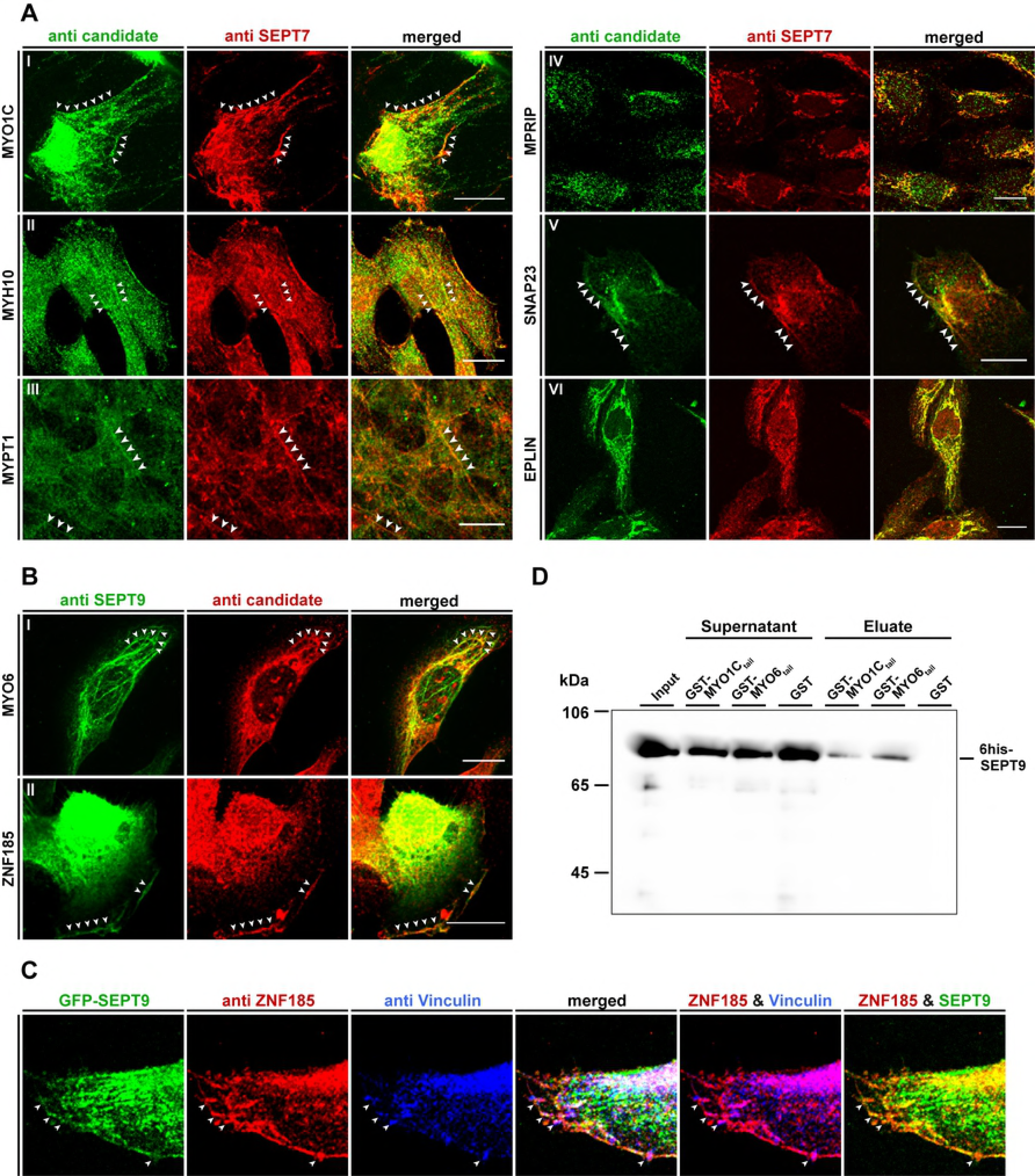
Colocalization of interaction partners with the septin cytoskeleton in 1306 fibroblast cells and Western blot analysis showing a direct interaction of SEPT9 with myosin motors. **A)** The indicated candidate proteins were immunostained with a suitable primary antibody, followed by an Alexa488 coupled secondary antibody. The endogenous septin cytoskeleton was immunostained by an anti-SEPT7 primary antibody followed by an Alexa555 coupled secondary antibody. White arrowheads mark colocalizing structures. **B)** Colocalization of MYO6 and ZNF185 with the endogenous septin cytoskeleton by immunostaining of SEPT9 with an anti-SEPT9 primary antibody followed by an Atto488 coupled secondary antibody. Due to species incompatibility of the available primary antibodies against MYO6 and ZNF185, these two candidates had to be visualized via an Alexa555 coupled secondary antibody. White arrowheads mark colocalizing structures. **C)** Colocalization of ZNF185, Vinculin and SEPT9 in GFP-SEPT9 expressing cells. ZNF185 was visualized as in figure part B and Vinculin was visualized by an antibody directly coupled to Alexa647. White arrowheads mark sites of focal adhesions. Images in parts A-C were assembled from z-projections. The scale bars represent 20 µM. **D)** *In vitro* binding experiment showing direct binding of 6his-SEPT9 to tail fragments of myosin motors. GST-MYO6_835-1294_, GST-MYO1C_809-1063_ and free GST as control were immobilized on Glutathione Sepharose and incubated with 2 µM 6his-SEPT9. Samples of the SEPT9 input, the supernatand after incubation and the eluate from the beads were subjected to SDS-PAGE and subsequent Western blot analysis using an anti-his primary antibody.

SNAP23 is part of the SNARE complex [37] and involved in docking and fusion of vesicles with their target membrane. It colocalized with SEPT9 at distinct, small dot-like structures at the cell periphery (Fig. 3A, VI and Fig. 4A, V). In human podocytes it is part of a complex containing SEPT7 and MYH9 [38]. MYH9, the heavy chain of non muscle myosin IIA, is not present in our data set and thus we cannot confirm such a complex in fibroblasts. Other septin subunits, namely SEPT5, SEPT7 and SEPT8, have been reported to participate in vesicle docking to the target membrane [38–40]. Through SEPT2, septins regulate exocytosis by dynamically interacting with components of the exocytosis apparatus [10]. A contribution of SEPT9 in vesicle transport and docking can be anticipated since other constituents of the vesicle trafficking machinery (VAMP2, VABP, COPA, SEC22B) were already identified in a high throughput MS screen as presumable SEPT9 interactors [41].

For two of the three LIM domain containing candidate proteins, EPLIN and ZNF185, we could show a colocalization with the septins in fibroblast cells. EPLIN localized with intracellular SEPT9 decorated filamentous structures (Fig. 3A, VII and Fig. 4A, VI), whereas ZNF185 colocalized exclusively with filaments at the cell cortex (Fig. 3A, VII and Fig. 4B, II). The used antibody for the third protein, LMO7, was unspecific and thus we employed a LMO7-GFP fusion protein to detect colocalization. In transiently transfected cells LMO7-GFP colocalized widely with the septin cytoskeleton (Fig. 3B). LMO7 is reported to bind the nuclear membrane protein EMERIN which is a key player in Emery–Dreifuss muscular dystrophy. Moreover, it shuttles between the plasma membrane and the nucleus [42] and seems to control mitosis progression and exerts an effect on the splindle assembly chekpoint [32]. LMO7 is also localized at focal adhesions [43]. Through the outcome of a high throughput interaction screen, LMO7 was linked to the actin cytoskeleton by its presumable interaction with different myosins (including MYO1C), anilin, beta-actin and EPLIN [44]. EPLIN in turn is a regulator of actin dynamics by bundling actin filaments [45] and provides a direct physical link between the actin cytoskeleton, the cadherin–catenin complex in adherens junctions [46] and the septins (see above). It interacts furthermore with CAV1 [47], one of our *bona fide* SEPT9 interactors that eluded from IF validation due to the unspecificity of the used antibody.

ZNF185 was previously found to interact with F-actin and to be enriched in focal adhesions [33]. Adherens junctions and focal adhesions are the two anchoring junction systems that have actin as cytoskeletal attachment system whereas desmosomes and hemisdesmosomes are attached via intermediate filaments. Adherens junctions and focal adhesions connect cells with other cells or with the extracellular matrix, respectively [48]. LMO7, EPLIN and ZNF185 serve as intracellular adaptor proteins in these cell junction systems.

Septins associate with the distal ends of radial stress fibers anchored to focal adhesions. It is supposed that septins mediate the anchoring of radial to transverse arc stress fibers. If this event occurs between fibers of opposing focal adhesions, septins may promote the generation of ventral stress fibres (summarized in [49]).

We stained ZNF185 and Vinculin, a focal adhesion resident protein, simultaneously in GFP-SEPT9 expressing cells (Fig. 4C). Indeed, ZNF185 colocalizes with SEPT9 decorated septin filaments as seen before and is enriched in focal adhesions. The focal adhesions seem to be connected by the septin decorated filaments but SEPT9 does not colocalize inside the adhesion complexes. This finding fosters the assumption that anchoring of the septins to the focal adhesion associated fibers is mediated by ZNF185.

Myosin phosphatase targeting subunit (MYPT1) is a non-catalytic subunit of myosin phosphatase and has a number of documented funtions in cytoskeleton organization, cell migration and cytokinesis [50]. It is located along stress fibers in fibroblasts and also to cell adhesions [51,52]. In a high throughput screen, MYPT1 was found as *bona fide* SEPT9 interactor [44]. In our cell culture model, MYPT1 colocalizes with SEPT9 decorated filaments spanning almost the whole length of the cell (Fig. 3A, IV and 4A, III).

Another protein interacting with stress fiber associated myosin phosphatase is MPRIP. This interaction is suggested to participate in the recruitment of myosin phosphatase to dephosphorylate myosin light chains on stress fibers [53]. As we found SEPT9 colocalizing with both MYPT1 and MPRIP (Fig. 3A, V and 4A, IV), we suggest that the septins serve in this context as a scaffold that bring myosin phosphatase, its regulatory subunits and its intracelllar substrates in proximity.

Apart from these myosin regulatory proteins, we could show that three myosins motors colocalized with the septin cytoskeleton, MYO1C, MYH10 and MYO6. All are unconventional, non-muscle myosins and are responsible for intracellular movements. Myosin I myosins (among these MYO1C and MYO1B) act as linkers between the actin cytoskeleton and membranes during exocytosis [54]. MYO1C was found to be required for the insulin stimulated, actin dependent translocation of GLUT4 containig vesicles towards the plasma membrane [55]. A similar context was reported for MYH9 in podocyte cells. Here, GLUT4 vesicle docking was found to be dependent of a complex built of MYH9, SNAP23 and SEPT7 [38]. As we have identified SNAP23 as SEPT9 interactor (see above), it can be anticipated that a similar complex consisting of MYO1C, SNAP23 and SEPT9 exists in fibroblasts. GLUT4 is expressed in dermal fibroblast cells [56]. Indeed, MYO1C colocalized with SEPT9 at the cell cortex (Fig. 4A, I and 5A, I).

MYO6 is an unconventional myosin that moves towards the minus end of actin filaments [35]. It colocalized with SEPT9 with dotted structures inside the cell or with bended filaments below the cortex (Fig. 3A, II and 4B, I). MYO6 is involved in targeted membrane transport during cytokinesis [36] and in the transport of uncoated endocytic vesicles from the actin rich cell periphery towards the endosome [57], a process that is also maintained by the septins. Septins were shown to regulate the formation of endocytic carriers at the plasma membrane and participate in endosomal sorting (reviewed in [58]). Possibly these functions are maintained by the interaction of SEPT9 with MYO6.

Are interactions of MYO6 and MYO1C with SEPT9 direct or mediated by adapter proteins? To answer this questions, we constructed GST fusion proteins of C-terminal tail fragments of these two myosins and performed a pulldown with recombinantly expressed, 6his tagged SEPT9. SEPT9 specifically interacts with both GST-MYO6_835-1294_ and GST-MYO1C_809-1063_ *in vitro* (Fig. 4D).

Another septin, SEPT2, is reported to interact directly with myosin motors, particularly with a tail fragment of non muscle myosin IIA which contains the myosin heavy chain MYH9 [34]. MYH9 was not identified as interactor of SEPT9 in our screen but the heavy chain of myosin IIB, MYH10, showed a colocalization with SEPT9 decorated filaments (Fig. 3A, III and 4A, II). We propose an isoform specific interaction with certain septin subunits -in this case myosin IIA with SEPT2 and myosin IIB with SEPT9. Based on the findings by Joo etc al, we propose also a direct interaction of MYH10 with SEPT9.

The interaction of SEPT2 with MYH9 was found to be important for the stability of stress fibers and loss of the interaction caused instability of the ingressed cleavage furrow in dividing cells [34]. Thus SEPT2 acts in this context as a functional unit rather than “only” a scaffold. The direct interaction of SEPT9 with myosin motors point towards a similar, functional, role of SEPT9 especially in vesicle docking during exo-and endocytosis. Our herein presented direct interactions of SEPT9 with myosin motors open thus new perspectives towards the role of the septins in intracellular processes that require further investigations.

## Materials and Methods

### Generation of the constructs

A pEGFP-N1 based expression plasmid for GFP-SEPT9_i1 was kindly provided by Elias Spiliotis (Drexel University, PA, USA). To generate a TAP-SEP9 construct, the GFP sequence was excised with the *Nhe*1 and *Bsr*G1 restriction sites and replaced by a sequence encoding for a N-terminal TAP tag which was amplified from the plasmid pBS1761 (Euroscarf) (primers ^5’^gcac<u>gctagc</u>atgataacttcgtatagcatac^3’^ and ^5’^gctg<u>tgtaca</u>gcttatcgtcatcatcaagtg^3’^; restriction sites underlined).

A mCherry-SEPT9 fusion portein was generated by replacing the GFP sequence by the mfCherry tag which was amplified from an expression plasmid kindly provided by Helge Ewers (FU Berlin, Germany).

To generate a 6his fusion protein for expression in *E.coli*, SEPT9 was PCR amplified (primers ^5’^ccgaa<u>ggccagcacggcc</u>gaaaacctgtacttccagggtaagaagtcttactcaggagg^3’^ and ^5’^ctgtg<u>ggccaaaaaggcc</u>ttatcacatctctggggcttctgg^3’^; restriction sites underlined) from the GFP expression plasmid and cloned into the in-house constructed expression plasmid pES allowing for the expression of N-terminal 6his fusion proteins in *E.coli*. An EGFP-MYO6 expression plasmid was kindly provided by Hans-Peter Wollscheid (IMB, Mainz, Germany). A C-terminal tail fragment of MYO6 spanning amino acids 835-1294 was PCR amplified (primers ^5’^gctag<u>ggatcc</u>aaacctcgcattgatggtctg^3’^ and ^5’^ctag<u>gaattc</u>cctactttaacagactctgcagc^3’^; restriction sites underlined) from this plasmid and cloned in the pGEX2T plasmid (GE Healthcare) yielding a N-terminal GST fusion protein for expression in *E. coli*. A C-terminal tail fragment of MYO1C spanning amino acids 809-1063 was PCR amplified (primers ^5’^gctag<u>ggatcc</u>ctggaccatgtgcgcacc^3’^ and ^5’^gctagtcgacgttcaccga<u>gaattc</u>agccgtg^3’^; restriction seites underlined) from cDNA obtained from in house prepared 1306 fibroblast mRNA and cloned in the pGEX2T plasmid.

A pcDNA-FRT/TO-GFP-LMO7 plasmid was obtained from the MRC Protein Phosphorylation and Ubiquitinylation Unit of the University Dundee, Ireland.

### Cell culture, immonofluorescence and microscopy

Immortalized 1306 skin fibroblast cells [59] were a kind gift of Sebastian Iben (Ulm University, Germany). The cells were cultivated in DMEM (Thermo Scientific) supplemented with 10% FCS (Thermo Scientific) under humidified 5 % CO_2_ atmosphere. For SILAC, TAP-SEPT9_i1 expressing cells were grown in SILAC-DMEM (Thermo Scientific) supplemented with 10 % dialyzed FBS, 0.398 mM L-Arginine-^13^C_6_ ^15^N_4_ hydrochloride (Sigma) and 0.798 mM L-Lysine-^13^C_6_ ^15^N_2_ (Sigma). Plasmids were transfected by standard calicium phosphate based transfection. For the generation of stable cell lines, Geneticin (G418, Formedium) was added 48 h post-transfection at a concentration of 500 ng/µl. After selection, cells were maintained under 250 ng/µl G418. The successful expression of the tagged SEPT9 constructs was verified by Western blotting and immunofluorescence (IF) using anti-Protein A (Sigma) and anti-SEPT9 (clone 2C6, Sigma) antibodies.

Cells for IF were grown on sterile coverslips placed in a 6-well cell culture plate in 2 ml medium or in 8-well tissue culture chamber slides (Sarstedt) previously coated with Poly-L-Lysine. The cells were fixed in 3 % Paraformaldehyde and permeabilized with 1 % TX-100 in DPBS (Biochrom) for 5 min at room temperature. After blocking with 10 % BSA, cells were incubated for 1 h at 37 °C with primary antibody followed by the respective secondary antibody. A list of all used antibodies for IF including the dilution is provided in Supplementary Table 2.

Cover slips were mounted with fluorescent mounting medium (Dako) and observed using an Observer SD confocal microscope (Zeiss) equipped with 488 nm, 547 nm and 647 nm diode lasers, a 63-fold Plan-Apochromat objective with lens aperture 1.4 and an Evolve 512K EMCCD camera (Photometrics). Image processing was done with the aquisition software of the microscope (Zen Vers. 2 blue, Zeiss) and with ImageJ.

### Affinity purification of SEPT9 complexes

Affinity purification of SEPT9 complexes was performed separately for TAP-SEPT9 and GFP-SEPT9 in a “mixing after purification” (MAP) approach.

Cells were grown to 80% confluency in one 175 cm^2^ bottle. The medium was aspirated, the cells washed once with PBS and subsequently detached using a cell scraper. The detached cells were pelleted and ice-cold lysis buffer (50 mM HEPES pH 7.5, 150 mM NaCl, 0.2 mM EDTA, 0.1% Triton X100, 8.3 μM Antipain, 0.3 μM Aprotinin, 1 mM Benzamidine, 1 μM Bestatin, 10 μM Chymostatin, 1.5 μM Pepstatin A, 5 μM Leupeptin, 1 mM PMSF, 10 mM β-Glycerophosphate, 10 mM NaF, 1 mM Sodium Orthovanadate) was added. The volume of lysis buffer including protease inhibitors was adjusted dependent on the cell pellet volume (approximately double pellet volume). For complete lysis, cells were incubated on ice and vortexed for 10 s at highest settings every 5 min for a total of 30 min followed by centrifugation at 13,000 × g for 10 min at 4 °C. The supernatant was directly used for separation by SDS-PAGE and Western blot analysis or affinity purification.

After cell lysis and determination of the protein concentration by a Bradford assay, the extract was adjusted to 10 % Glycerol and the concentrations of the extracts were balanced by dilution of the higher concentrated extract with buffer. CNBr activated Sepharose (GE Healthcare) was coupled with human IgG (MP Biomedicals) according to the manufacturer’s protocol and stored as slurry in PBS/10 mM NaN_3_. The resulting hsIgG-Sepharose was equilibrated with lysis buffer and added to the cell lysates (15 µl slurry per mg of protein). Incubation was performed by rotating with 5 rpm overnight at 4 °C in an overhead shaker. The following day, an equivalent volume of hsIgG-Sepharose was added and incubated for additional 2 h at 4 °C while rotating at 5 rpm. The sepharose was pelleted at 100 ×g for 3 min at 4 °C and transferred into Mobicol “F” columns (Mobitec) equipped with a 35 µm filter. After washing the slurry with 50-fold volume TAP buffer (20 mM Tris pH 7.5, 80 mM NaCl, 10 % Glycerol, 1 mM DTT, 8.3 μM Antipain, 1 μM Bestatin, 10 μM Chymostatin, 1.5 μM Pepstatin A, 5 μM Leupeptin, 1 mM PMSF, 10 mM β-Glycerophosphate, 1 mM Sodium Orthovanadate), the columns were locked and two slurry volumes of TAP buffer was added. Proteins were eluted from the matrix by boiling in 2× Laemmli-buffer for 2 min at 95 °C and equal volumes of both sample and control were mixed for the MAP approach. The samples were jointly separated by SDS-PAGE on 6-12 % BOLT Bis-Tris gradient gels (Thermo Scientific). The gel was stained by colloidal coomassie staining by pre-incubating in staining solution (34 % (v/v) methanol, 2 % (v/v) phosphoric acid, 17 % (w/v) ammonium sulfate) for 1 h at RT and subsequent addition of 0.066 % (w/v) Coomassie brilliant blue G-250. After incubation overnight, the gel was de-stained with ddH_2_O and subjected to MS analysis. Affinity purification was performed as triplicate.

### Mass spectrometry

Each SDS-PAGE gel was cut into 14 pieces. Individual pieces were washed thrice by alternating incubation in 50 mM ammonium bicarbonate and 25mM ammonium bicarbonate / 50% Acetonitrile (ACN) for 10 minutes each. Following vacuum drying, samples were reduced with 5 mM DTT for 20 min at RT and subsequently alkylated with iodoacetamide for 20 min at 37 °C. After a second vacuum drying step, proteins were subjected to tryptic digest overnight at 37 °C as described elsewhere [60]. Peptides were extracted in two rounds by adding 20 µl 0.1 % Trifluoroacetic acid (TFA) / 50 % ACN and incubation in an ultrasonic bath for 10 min each. ACN was evaporated and samples filled to 15 µl with 0.1% TFA.

Samples were measured using an LTQ Orbitrap Velos Pro system (Thermo Fisher Scientific) online coupled to an U3000 RSLCnano (Thermo Fisher Scientific) as described previously [61], with the following modifications: Separation was carried out using a binary solvent gradient consisting of solvent A (0.1 % FA) and solvent B (86 % ACN, 0.1 % FA). The column was initially equilibrated in 5 % B. In a first elution step, the percentage of B was raised from 5 % to 15 % in 5 min, followed by an increase from 15 % to 40 % B in 30 min. The column was washed with 95 % B for 4 min and re-equilibrated with 5 % B for 25 min.

Database search was performed using MaxQuant Vers. 1.5.2.8 (www.maxquant.org) [26]. For peptide identification, MS/MS spectra were correlated with the UniProt human reference proteome set (www.uniprot.org) supplemented with the sequence of the TAP-SEPT9 construct, employing the build-in Andromeda search engine [62]. The respective SILAC amino acids were selected, while Carbamidomethylated cysteine was considered as a fixed modification along with oxidation (M), and acetylated protein N-termini as variable modifications. False discovery rates were set on both, peptide and protein level, to 0.01. MaxQuant normalized ratios were used for statistical evaluation. If no ratio was calculated by MaxQuant, ratios were calculated manually in case overall intensity exceeded 1E6 to allow for quantitation of IP-specific proteins. For statistical evaluation, Significance B was calculated using Perseus 1.5.0.15 (www.maxquant.org) with default parameters.

The complete data evaluation results including statistics is provided in Supplementary Table 1.

### In vitro binding assays

6his-tagged SEPT9 was expressed in 250 ml LB medium at 37 °C for 3 h after addition of 0.1 mM IPTG at an OD_6oo_ of 1.0. The cell pellet was resuspended in 25 ml IMAC A (50 mM KH_2_PO_4_ pH 7.5, 300 mM NaCl, 20 mM Imidazole) containing e-complete protease inhibitor cocktail (Roche) and lyzed by addition of lysozyme and ultrasound treatment.

The crude extract was subjected to IMAC chromatography using a 5 ml HisTrap excel column (GE Healthcare) and an ÄKTA Purifier chromatography system (GE HEalthcare). The protein was eluted from the column in three consecutive elution steps of 15 %, 40 % and 100% IMAC B (50 mM KH_2_PO_4_ pH 8.0, 300 mM NaCl, 200 mM Imidazole). We considered approx. 70 % purity after IMAC (judged by eye from a SDS-PAGE gel) as sufficient for further qualitative binding experiments.

The protein was transferred into PBS using a PD10 desalting column (GE Healthcare), concentrated and immediately used for binding assays.

GST-MYO6_835-1294_ and GST-MYO1C_809-1063_ were expressed in 50 ml SB medium at 18 °C for 20 h after addition of 1.0 mM IPTG at an OD_6oo_ of 1.0. Cell lysis was performed as described above in PBS and the GST fusion proteins were immbilized onto each 100 µl Glutathione Sepharose (GE Healthcare) from the crude extract. The beads were incubated with 2 µM 6his-SEPT9 for 1h and bound protein was detected after washing and elution from the beads by Western blot analysis using a mouse anti-6his primary antibody (Sigma).

### Data availability and supplementary information

The mass spectrometry proteomics data have been deposited to the ProteomeXchange Consortium via the PRIDE [63] partner repository with the dataset identifier PXD010773.

An Excel sheet containg the complete data evaluation results including statistics is provided as Supplementary Table 1.

IFs of not colocalizing candidate proteins are provided as Supplementary Figure 1. A referenced list of all used antibodies is provided as Supplementary Table 2.

## Acknowledgements

Matthias Hecht is supported by the International Graduate School in Molecular Medicine (Ulm University, Ulm, Germany).

Reinhild Rösler is supported by the SFB grant “1074-experimental models and clinical translation in leukemia” of the DFG.

## Author contributions

Matthias Hecht performed the experiments and prepared the figures. Reinhild Rösler and Sebastian Wiese conducted the MS measurements and MS data evaluation. Nils Johnsson improved the manuscript and helped to analyze the data. Thomas Gronemeyer performed the *in vitro* binding experiments, conceived the study and wrote the manuscript.

---

**Supplementary Figure 1. IF of candidate proteins not colocalizing with the septin cytoskeleton. A)** IF experiments in GFP-SEPT9 expressing cells. The indicated candidate proteins were immunostained with a suitable primary antibody, followed by an Alexa555 coupled secondary antibody. GFP-SEPT9 was observed directly via its GFP. **B)** IF experiments in 1306 wildtye cells. Indicated candidate proteins were immunostained with a suitable primary antibody, followed by an Alexa488 coupled secondary antibody. The endogenous septin cytoskeleton was immunostained by an anti-SEPT7 primary antibody followed by an Alexa555 coupled secondary antibody. Images were assembled from z-projections. The scale bars represent 20 µM.

